# Ligand Binding Free Energy Evaluation by Monte Carlo Recursion

**DOI:** 10.1101/2022.08.01.502202

**Authors:** Joao Victor de Souza, Victor H. R. Nogueira, Alessandro S. Nascimento

## Abstract

The correct evaluation of ligand binding free energies by computational methods is still a very challenging active area of research. The most employed methods for these calculations can be roughly classified into four groups: (*i*) the fastest and less accurate methods, such as molecular docking, designed to sample a large number of molecules and rapidly rank them according to the potential binding energy; (*ii*) the second class of methods use a thermodynamic ensemble, typically generated by molecular dynamics, to analyze the endpoints of the thermodynamic cycle for binding and extract differences, in the so-called ‘end-point’ methods; (*iii*) the third class of methods is based on the Zwanzig relationship and computes the free energy difference after a chemical change of the system (alchemical methods); and (*iv*) methods based on biased simulations, such as metadynamics, for example. These methods require increased computational power and as expected, result in increased accuracy for the determination of the strength of binding. Here, we describe an intermediate approach, based on the Monte Carlo Recursion (MCR) method first developed by Harold Scheraga. In this method, the system is sampled at increasing effective temperatures, and the free energy of the system is assessed from a series of terms *W*(*b, T*), computed from Monte Carlo (MC) averages at each iteration. We show the application of the MCR for ligand binding with datasets of guest-hosts systems (N=75) and we observed that a good correlation is obtained between experimental data and the binding energies computed with MCR. We also compared the experimental data with an end-point calculation from equilibrium Monte Carlo calculations that allowed us to conclude that the lower-energy (lower-temperature) terms in the calculation are the most relevant to the estimation of the binding energies, resulting in similar correlations between MCR and MC data and the experimental values. On the other hand, the MCR method provides a reasonable view of the binding energy funnel, with possible connections with the ligand binding kinetics, as well. The codes developed for this analysis are publicly available on GitHub as a part of the `LiBELa`/`MCLiBELa` project (https://github.com/alessandronascimento/LiBELa).

## 1. Introduction

Molecular interactions are central to most biological processes. Cell signaling depends on the interaction of receptors and hormones (e.g.[1]); metabolism and catabolism depend on enzyme-substrate recognition and many proteins exert their biological functions by forming dimers or higher-order oligomers. Despite the pivotal relevance of molecular interactions, the accurate evaluation of the strength of binding involving two molecular species in a biological context is still a challenge[2].

The most relevant thermodynamic quantity associated with the binding is the change in free energy. This quantity can be rigorously computed from molecular dynamics simulations using the alchemical methods of free energy perturbation (FEP) or thermodynamic integration (TI)[3]. Other computer inexpensive methods have also been proposed, based on the end-points of the thermodynamic cycle[4,5], or a fast evaluation of the partition function based on limited sampling[6,7]. In the best scenarios, binding free energies can be computed with a precision of around 1 kcal/mol, for the alchemical methods, at the cost of several (typically dozens of) simulations[8]. Another approach for accurate computation of the ligand binding free energy involves the simulation of an unbinding process, using the funnel metadynamics[9], for example. In this case, appropriate collective variables must be chosen, and the unbinding should be properly sampled in MD simulations[10]. Finally, a different approach focuses on the direct estimation of the free energy using a Markov State Modeling (MSM) of the *macrostates* bound and unbound from several rapid MD or MC simulations. Guallar and coworkers showed promising results using MC simulations in `PELE` combined with MSM for the estimation of ligand binding free energies[11,12]. An emerging area of intense research combines the alchemical method FEP with machine learning, allowing better estimation of parameters and better precision in free energy estimation[13,14], at the cost of losing throughput in calculations.

Following classical thermodynamics, the change in free energy due to binding is given by the sum of the change in macroscopic energy (ΔU) and the change in the entropy of the system at a given temperature (-TΔS). Although conceptually simple, the evaluation of this thermodynamic quantity involves several challenges. The macroscopic energy *U* can be assumed to be the average energy for the different microscopic states sampled by molecular dynamics (MD) or Monte Carlo (MC) simulations[15], given that a good sampling of thermally accessible conformations is achieved. The estimation of entropy, on the other hand, is less straightforward. As pointed out by Edholm and Berendsen[16], the multidimensional distribution of variables of the system hinders an accurate evaluation of changes in the system entropy. In addition to the problem of evaluating the thermodynamic quantities associated with binding, the sampling of thermally accessible and relevant microstates is itself an important issue to be addressed by free energy calculation methods[17].

Following the concept of the folding landscape, as proposed by Onuchic and Wolynes[18–20], one could generalize the concept to the context of ligand binding. In this context, the ligand binding process can be thought of as a free energy funnel as a function of the ligand-receptor coordinates during the binding process. The depth of the funnel is related to the energy change (ΔU) due to binding whereas the funnel radius is related to the change in entropy due to ligand binding. These ideas have already been proposed[21–23], however, some difficulties remain in the accurate estimation of the binding energies. Although the advances in GPU computing allow a better sampling of the phase space coupled with better force fields and a myriad of simulation methods, the small differences in binding energies, as compared to the absolute energies for the bound complex and its isolated parts, make the precise computation of the binding energies still a challenge.

Here, we evaluated a fast estimation of the binding free energy for a set of more than 70 host-guest systems using the Monte Carlo Recursion (MCR) approach, first proposed by Harold Scheraga[24]. in this work, we aimed to use computationally quick methods without compromising the accuracy, which could be used to evaluate binding energies in lower-end workstations. Our tests showed that MCR obtains a good correlation with experimental data and is fast to be used in large-scale drug discovery campaigns.

## 2. Methods

The methods in the sections below describe the implementation of a Monte Carlo Recursion calculation designed to provide a better ranking of ligands according to their binding free energy. The implementation is devised in a scenario where ligands identified in docking campaigns, for example, can be prioritized with a better calculation of energy scores with MCR than with the docking scores. In this context, we assume that the receptor can be treated as a rigid receptor and the solvent effects can be modeled using an empirical implicit solvation model.

### 2.1 Test Sets

To evaluate the interaction energies computed from MC simulations, a set of 30 receptor-ligand complexes were selected. The complexes are experimental crystallographic structures of the T4 lysozyme (T4L) mutant L99A and the double mutants L99A/M102Q[25] and L99A/M102E bound to several small ligands (or fragments), for which the experimental binding free energies are available in the literature and compiled in the database *BindingDB*[26] and from Mobley and Gilson’s[27] work. The PDB IDs for these complexes are given in Table 1, together with their experimental binding free energy.

**Table 1.** Experimental and computed binding free energies for the T4L dataset. Experimental errors, when available, are also reported. The computed binding energy error refers to the standard deviation computed for N=10 simulations for each complex. The experimental data were taken from the references shown in the last column.

| PDB | Ligand | Mutation | Experimental Free Energy (kcal/mol) | Computed Binding Energy (kcal/mol) | Computed Binding Energy Error (kcal/mol) | Reference |
| --- | --- | --- | --- | --- | --- | --- |
| 182L | Benzofuran | L99A | -5.46 ± 0.03 | -12.5 | 0.2 | [46] |
| 183L | Indene | L99A | -5.13 ± 0.01 | -13.4 | 0.1 | [46] |
| 184L | Isobutylbenzene | L99A | -6.51 ± 0.06 | -13.0 | 0.3 | [46] |
| 185L | Indole | L99A | -4.89 ± 0.06 | -12.8 | 0.4 | [46] |
| 186L | N-butylbenzene | L99A | -6.70 ± 0.02 | -14.79 | 0.06 | [46] |
| 187L | *p*-Xylene | L99A | -4.67 ± 0.06 | -11.0 | 0.2 | [46] |
| 188L | *o*-Xylene | L99A | -4.60 ± 0.06 | -10.7 | 0.2 | [46] |
| 1LGW | 2-Fluoroaniline | L99A/M102Q | -5.53 | -9.1 | 0.2 | [25] |
| 4W52 | Benzene | L99A | -5.19 ± 0.16 | -8.7 | 0.2 | [47] |
| 4W53 | Toluene | L99A | -5.52 ± 0.04 | -10.30 | 0.01 | [47] |
| 4W55 | Propylbenzene | L99A | -6.55 ± 0.02 | -13.1 | 0.2 | [47] |
| 4W57 | N-butylbenzene | L99A | -6.70 ± 0.02 | -13.7 | 0.4 | [47] |
| 1LI2 | Phenol | L99A/M102Q | -5.24 | -9.2 | 0.2 | [25] |
| 1LI3 | 3-Chlorophenol | L99A/M102Q | -5.51 | -10.5 | 0.3 | [25] |
| 1LI6 | 5-methylpyrrole | L99A/M102Q | -5.25 | -8.3 | 0.1 | [25] |
| 1NHB | Phenylethane | L99A | -5.76 ± 0.07 | -10.8 | 0.1 | [46] |
| 3DMX | Benzene | L99A | -5.19 ± 0.16 | -8.6 | 0.3 | [48] |
| 3HT8 | 5-Chloro-2-methylphenol | L99A/M102Q | -5.42 | -12.01 | 0.04 | [49] |
| 3HTB | 2-Propylphenol | L99A/M102Q |  | -14.5 | 0.2 |  |
| 1XEP | Catechol | L99A/M102Q | -4.16 | -10.84 | 0.02 | [50] |
| 2RBO | 2-Nitrothiophene | L99A/M102Q |  | -12.38 | 0.03 |  |
| 3HU8 | 2-Ethoxyphenol | L99A/M102Q | -4.02 | -14.0 | 0.1 | [49] |
| 3HUK | Benzyl acetate | L99A/M102Q | -4.48 | -4.5 | 1.1 |  |
| 3HUA | 4,5,6,7-tetrahydro-1H-indole | L99A/M102Q | -4.61 | -14.39 | 0.03 | [49] |
| 3GUJ | Benzene | L99A/M102E | -5.8 | -9.2 | 0.3 | [51] |
| 3GUK | Toluene | L99A/M102E | -5.3 | -10.70 | 0.05 | [51] |
| 3GUL | Ethylbenzene | L99A/M102E | -5.5 | -11.21 | 0.05 | [51] |
| 3GUM | *p*-Xylene | L99A/M102E | -4.9 | -11.1 | 0.2 | [51] |
| 3GUN | Aniline | L99A/M102E | -4.0 | -9.9 | 0.1 | [51] |
| 3GUO | Phenol | L99A/M102E | -5.0 | -10.89 | 0.07 | [51] |

In all cases, the receptor structure was parametrized with the AMBER FF14SB force field (atomic charges and Lennard-Jones parameters) using the `DockPrep` tool as available in UCSF Chimera[28]. The structural water molecules within 5 Å of the ligand binding site were maintained in the receptor structure. The ligands were parametrized using the General AMBER force field (GAFF)[29] version 2 using `ANTECHAMBER`[30] and AM1-BCC as the charge model.

Other datasets evaluated here include the cyclodextrin (CD) dataset, the curcubit[7]uril (CB7) dataset, and the BRD4 (first bromodomain of the BRD4 protein) dataset. For these datasets, the SYBYL MOL2 files were used as provided by David Mobley’s group[31]. For the CD and CB7 datasets, the atomic charges were computed using the RESP procedure[32] using `ANTECHAMBER`[30], fitting the electrostatic potential grids computed during the energy minimization in vacuo using HF/6-31G(d). For BRD4, the receptor protein was parametrized using UCSF Chimera[28] tool `DockPrep` with AMBER FF14SB[33] atomic charges.

### 2.2 Monte Carlo Implementation

The MC simulations were performed in a modified version of our ligand docking software named `LiBELA` (Ligand Binding Energy Landscape)[34]. The algorithm involves a random translation and rotation of the ligand within the receptor active site, followed by a random shift in the angle of each rotatable bond (RB). This set of 6 + 3*n*_*RB*_ variables defines a *move*. The energy for the new coordinates is computed and tested according to the Metropolis criterion. The random displacements for Monte Carlo moves are sampled using the random number generator based on the RANLUX2 algorithm as implemented in the GNU Scientific Library[35].

Before each *production* simulation, the system was equilibrated during 10^6^ MC accepted moves or steps, and, in the case of equilibrium MC simulations, a production simulation was conducted for 5×10^7^ MC steps. The maximal translation step was set to 0.5 Å for each direction, while the maximal rotation step and torsion scan step were set to 1.25 degrees. This modified version of `LiBELa` will be referred to hereafter as `MCLiBELa`. In order to speed up the calculations, the binding potentials are pre-computed in tridimensional grids[36,37] assuming a rigid receptor, as usual in docking calculations. Briefly, the Lennard-Jones attractive and repulsive terms due to the receptor atoms are pre-computed in a 40 × 40 × 40 Å cube with grid points spaced by 0.4 Å. The electrostatic potential is also stored in the grids together with the solvation term associated with the fragmental volume of the receptor atoms with a Gaussian distance-dependent weighting term (see below).

For the purposes of this work, a modified version of the original interaction energy model was adopted. Here, the interaction energy was evaluated as:

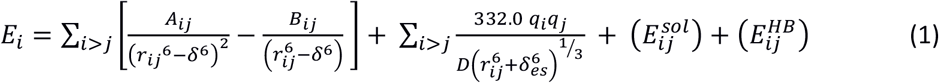

In this model, the first term described a softcore Lennard-Jones potential[38] between the ligand atoms (*i*) and receptor atoms (*j*), with a smoothing parameter δ set to 2.5 Å. The second term models the polar interactions, modeled as a smoothed Coulomb model. In this case, the dielectric ‘constant’ was set to interatomic distance, i.e., *D* = *r*_*ij*_, while the smoothing parameter δ_es_ was set to 2.5 Å. The third term, E_ij_^sol^, is a desolvation term, previously described by Stouten[39] and Verkivker[38] and also evaluated before in the context of ligand docking[40]. In this model, the affinity of each atom *i* to the polar solvent, *S*_*i*_, is modeled as a linear function of the square of the atomic charge:

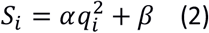

This empirical affinity is then combined with the volume of the solvent displaced upon interaction (solvent excluded volume), with a Gaussian envelope:

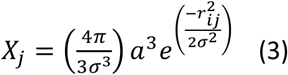

where *a* is the atomic radius of the atom *j* interacting with atom *i* and desolvating it, *r*_*ij*_ is the interatomic distance and σ is a constant (*σ*=3.5 Å). The final energy associated with ligand and receptor desolvation is given by[38]:

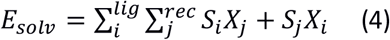

In this work, the parameter α was set to 0.1 kcal/(mol. e^2^) and β was set to -0.005 kcal/mol[40]. The last term in equation (1), E_ij_^HB^, explicitly models the hydrogen bonds through a 10-12 potential and a directionality term. The parameters are set to result in an interaction energy of -5.0 kcal/mol for a typical hydrogen bond and the directionality term *cos*^4^(*θ*), where θ is the angle between donor, hydrogen atom, and acceptor is used to ensure the proper geometry of the interaction.

For each ligand conformation being sampled, the ligand internal energy (E_lig_) is computed using the OpenBabel API[41] and the GAFF force field. The final energy (E_t_), as computed in this work, is the sum of the interaction energy (E_i_, in equation 2) and ligand internal energy:

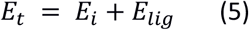

E_t_, as defined in equation (5), is the term used in the Metropolis criteria in the MC simulations.

When necessary, binding energies were computed in end-point calculations from equilibrium Monte Carlo data. For this purpose, MC simulations for the receptor-ligand complex and for the free ligand were used to compute average energies and their difference Δ*U*. The change in entropy was estimated by assuming that each configurational degree of freedom for the ligand (rotational, translational, and torsional) can be considered independent of each other in a first-order approach[42]. The entropy associated with each degree of freedom is computed as a Shannon entropy and the probabilities are computed by binning the observed values. The entropy differences are computed by subtracting the total entropy for the free ligand from the entropy for the bound ligand.

### 2.3 Monte Carlo Recursion

The Monte Carlo Recursion (MCR) method is rooted in the computation of the quantity *W*(*b,T*), defined as[24,43]:

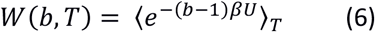

where b is a constant (*b*>1) and the brackets represent a Monte Carlo average, evaluated at an MC temperature T. Li and Scheraga noted that *W(b,T)* is connected to the system partition function *Q(T)* by [24,43]:

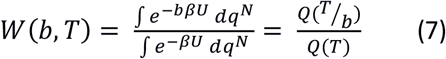

By iterating *b*, i.e., computing *W(b,T), W(b*_*2*_, *bb*_*2*_*T), W(b*_*3*_, *bb*_*2*_*b*_*3*_*T)*, …, etc, one arrives at:

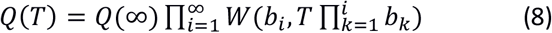

Using *Q*(∞) = *V*^*N*^, the configurational Helmholtz free energy *A*(*T*) can thus be given by:

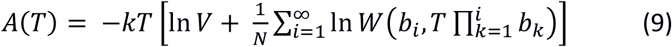

where 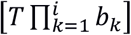 is an ‘*effective temperature*’ assigned to each iteration *i*. Although the free energy is computed from equation (9) from an infinite series, the terms in the summation approach zero as the temperature increases, conferring convergence to the method after a few recursion terms. For the results shown here, 12 terms were included in the calculations to ensure convergence, with coefficients *b*_*i*_ set to 1.5, 1.5, 2.0, 2.0, 2.0, 4.0, 8.0, 16.0, 32.0, 64.0, 128.0 and 256.0. For each term of the recursion, the system was allowed to equilibrate for 10^6^ MC steps, followed by 20 million MC steps of production simulation, used to calculate *W*(*b, T*).

The binding free energy is computed using the same simulation setup for the ligand sampled within the receptor structure (*A*_*complex*_) and for the free ligand (*A*_*ligand*_) and taking their difference: Δ*A*_*bind*_ = *A*_*complex*_ − *A*_*ligand*_. Ten independent simulations were set up for each system using different random seeds. The errors in the estimation of the binding energy were computed as the standard deviation among the ten simulations.

## 3. Results

The estimation of the binding energy involved in ligand-receptor recognition is of paramount relevance in the structure-based design of drugs. Despite its relevance, this estimation still represents a challenge for computational structural biologists. Shoichet’s and Irwin’s groups showed recently that the energy scores computed from docking calculations show only a modest correlation with ligand affinity[44,45], suggesting that the prioritization of a given compound after a virtual screening is still a challenge, especially if the project budget limits the number of compounds to be experimentally tested.

On the other hand, the most accurate methods for affinity estimation require expensive MD/MC simulations, making them impractical for the prioritization of a large number of compounds identified in a large to ultra-large docking campaign involving more than a hundred million compounds screened[44]. In this scenario, we looked for a methodology that could be accurate enough to suggest the best binders after a reasonable docking pose has been found but preserving a relatively low computational cost. In this context, we evaluated a Monte Carlo-based sampling method that could be able to refine the docking poses, but also ensure better linearity among the computed binding energies and the actual binding free energy.

After preliminary tests, the MC temperature was set to room temperature, i.e., 300 K, for all the simulations, ensuring that the ligand sampled conformations within the active site and close to the crystallographic pose. Taking the average over the 30 for T4 lysozyme complexes, each calculation took about 23 hours for complex and free ligand, running one MCR simulation per processor thread of Intel Xeon E5645 processors. Thus, the computational costs involved in the calculations are affordable with small in-house computing resources, even for the screening of hundreds of molecules, looking forward to the prioritization of compounds previously selected in ligand docking campaigns.

From a sampling point of view, the MCR method can be seen as a sort of ‘*inverse simulated annealing*’, in the sense that the effective temperature is constantly increased to sample high-energy states and finally resulting in convergence in the cumulative quantity In(*W*). This quantity will be dominated by the low energy terms, sampled in the first iterations, making it relevant to choose appropriate temperatures to start the simulations, as well as a reasonable set of parameters *b*_*i*_ for the MC recursion.

The convergence of the calculations of the binding energies was first analyzed. To assess whether the sampling obtained in the MCR simulations was appropriate and reached an equilibrium state, we analyzed the cumulative quantity In (*W*), for the complex, i.e., bound ligand and receptor (semitransparent blue points in Figure 1), as well as for the isolated ligand (semitransparent red points) and their difference (green points). As shown in Figure 1, the calculations reach a converged state with less than 10 iterations. The estimation of the free energies, as defined in equation (9), also revealed a convergence of this quantity to less than 0.1 kcal/mol after the 12 iterations in MCR.

**Figure 1.**
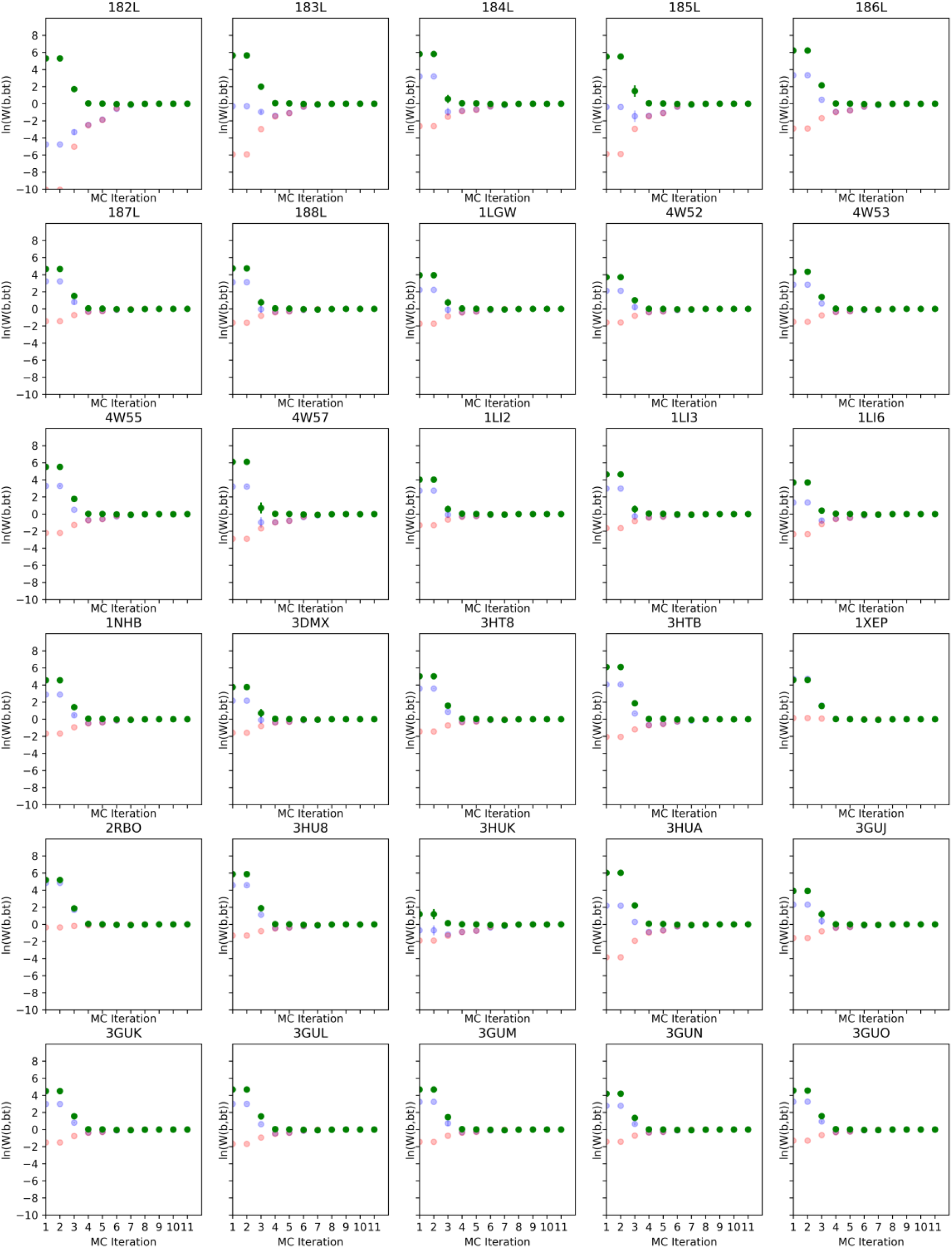
Convergence analysis of the MCR simulations for the T4L complexes. Each plot shows the cumulative quantity ln(W) for the complex (semitransparent blue points), for the ligand (semitransparent red points), and their difference (green points). Each point is represented as an average of 10 simulations with its error bars showing the standard deviation.

A closer inspection of the results shown in Figure 1 also shows the relevance of the first five terms in the recursion. The difference between the ligand-bound simulation and free ligand simulation resides in these first terms, where the ligand is sampling low-energy conformations. By reaching higher effective temperatures, the ligand is usually dissociated from the receptor and the sampling is mostly restricted to a free ligand in solution. In this scenario, the quantity In (*W*) is mostly the same for both simulations and tends to zero as the temperature increases.

This analysis is also shown for representative complexes of the T4L dataset in Figure 2. Here, the ligand-enzyme complexes 186L, 182L, and 184L are shown during the four first iterations (Figure 2, left). The panel shows the ligand sampling different conformations mostly within the binding pocket while the right panel shows the sampled conformations during the fifth iteration. The ligand is usually displaced in the fourth iteration. At the beginning of this iteration, the ligand is still found in the ligand pocket, but is rapidly displaced and starts to sample different conformations outside the ligand pocket and around the enzyme (Figure 2, left panel). The results shown here are in line with the values computed for the cumulative In(*W*) shown in Figure 1, where the differences between ligand-bound and free ligand simulations tend to zero after the fourth iteration.

**Figure 2.**
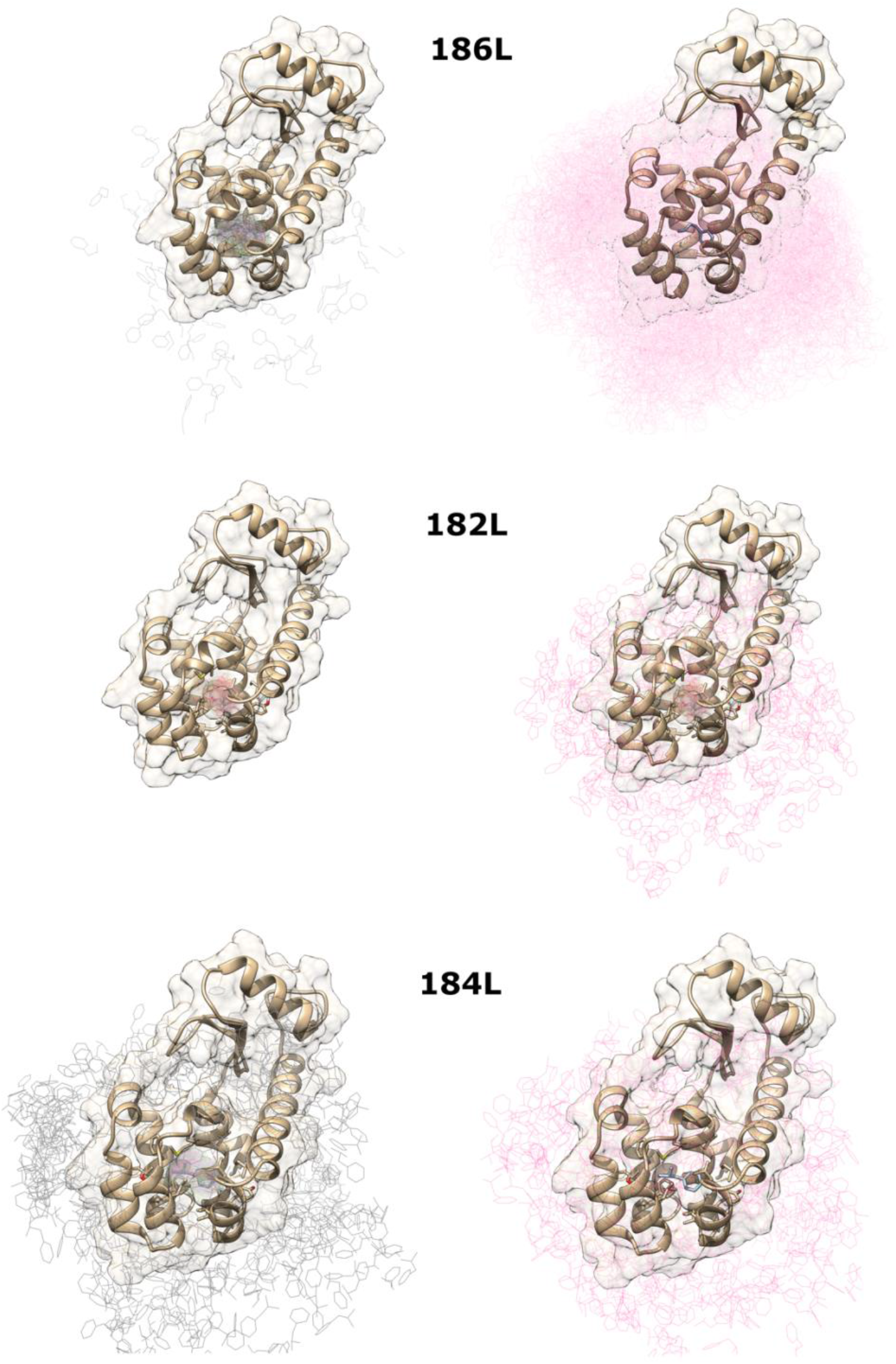
Coordinates for the ligand during an MCR simulation. Here, the systems 186L, 182L, and 184L are shown as representatives of the MCR simulations. (left) Most conformations sampled by the ligands are in the active site for iterations 1-4. In the 186L system (top) and 184L system (bottom), the ligand displacement is observed in the 4^th^ iteration. (right) The 5^th^ iteration of the MCR simulations is shown for the three ligand-receptor systems. The ligand is mostly unbound from the enzyme’s active site. The ligand conformations are shown in semitransparent wires.

The binding energies were computed following equation (9). For this calculation, the cubic volume sampled by the ligand was estimated by the maximal displacement in the center of mass in each direction for each iteration. Figure 3 shows the free energy computed for each ligand of the T4L dataset as a function of the MCR iteration. It can be observed, from the figure, that the computed binding free energies converge with less than 10 iterations.

**Figure 3.**
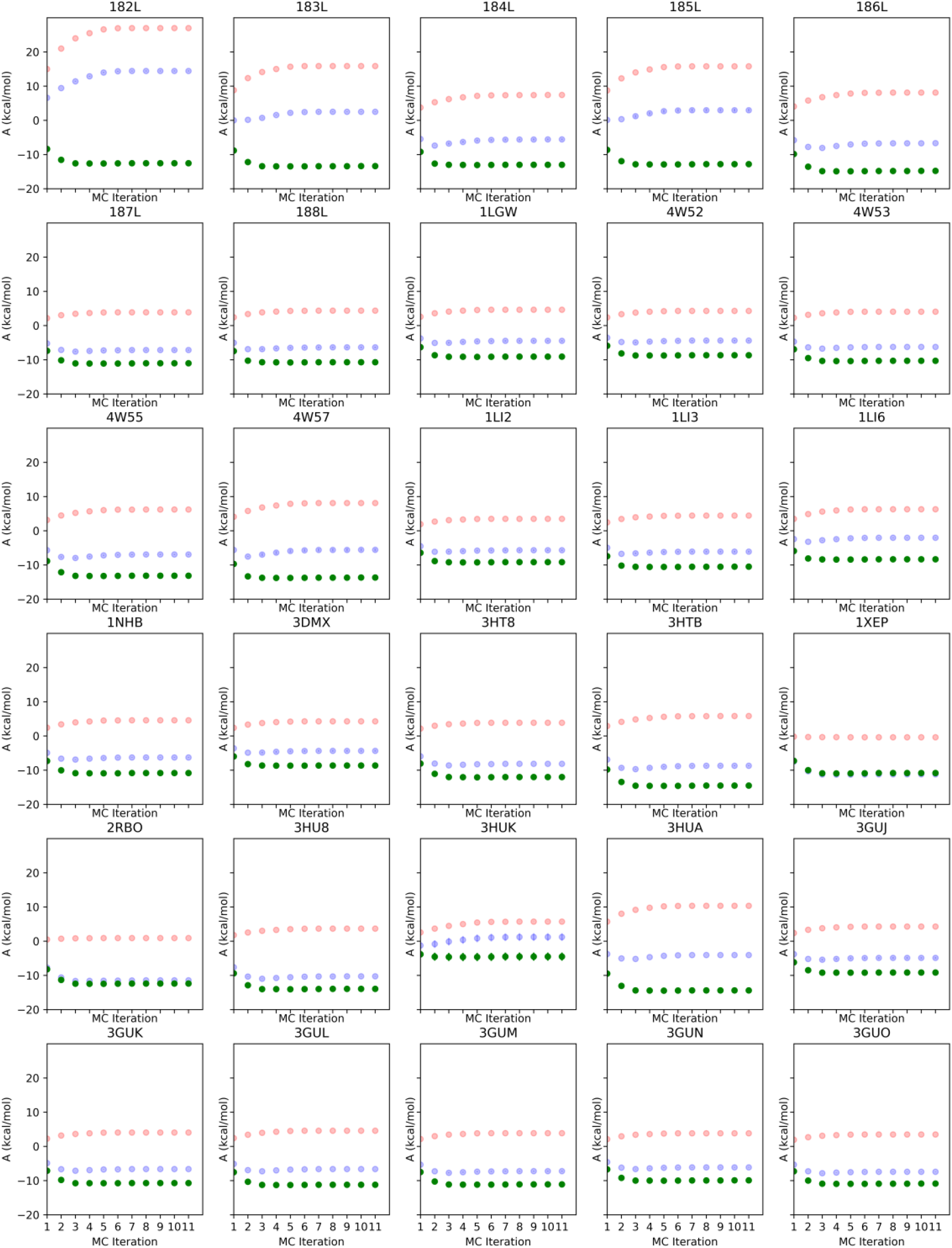
Free energy analysis of the MCR simulations for the T4L complexes. Each plot shows the Helmholtz free energy computed for the complex (semitransparent blue points), for the ligand (semitransparent red points), and their difference (green points). Each point is represented as an average of 10 simulations with its error bars showing the standard deviation.

A comparison between the computed energies and the experimental binding free energies showed only a modest correlation between experimental and computed binding data, as shown in Figure 4 (inset) and Table 1. Here some interesting issues must be highlighted. First, the computed energies are in the interval between -8 to -18 kcal/mol, while the experimental binding energies are found in the range of -3 to -7 kcal/mol, indicating that the MCR method, as applied here, significantly overestimates the binding energies. Given the assumptions made, which include a rigid receptor, a smoothed forcefield-based potential, and the computation of binding energies using the grid approach, one would not expect a precise estimation of the binding free energies, but a good correlation between the MCR computed energies and the experimental binding free energies. Interestingly, however, Pearson’s correlation between the binding free energies and the computed binding energies is reasonably weak for this dataset (*r* = 0.3), as shown in Figure 4 (inset).

**Figure 4.**
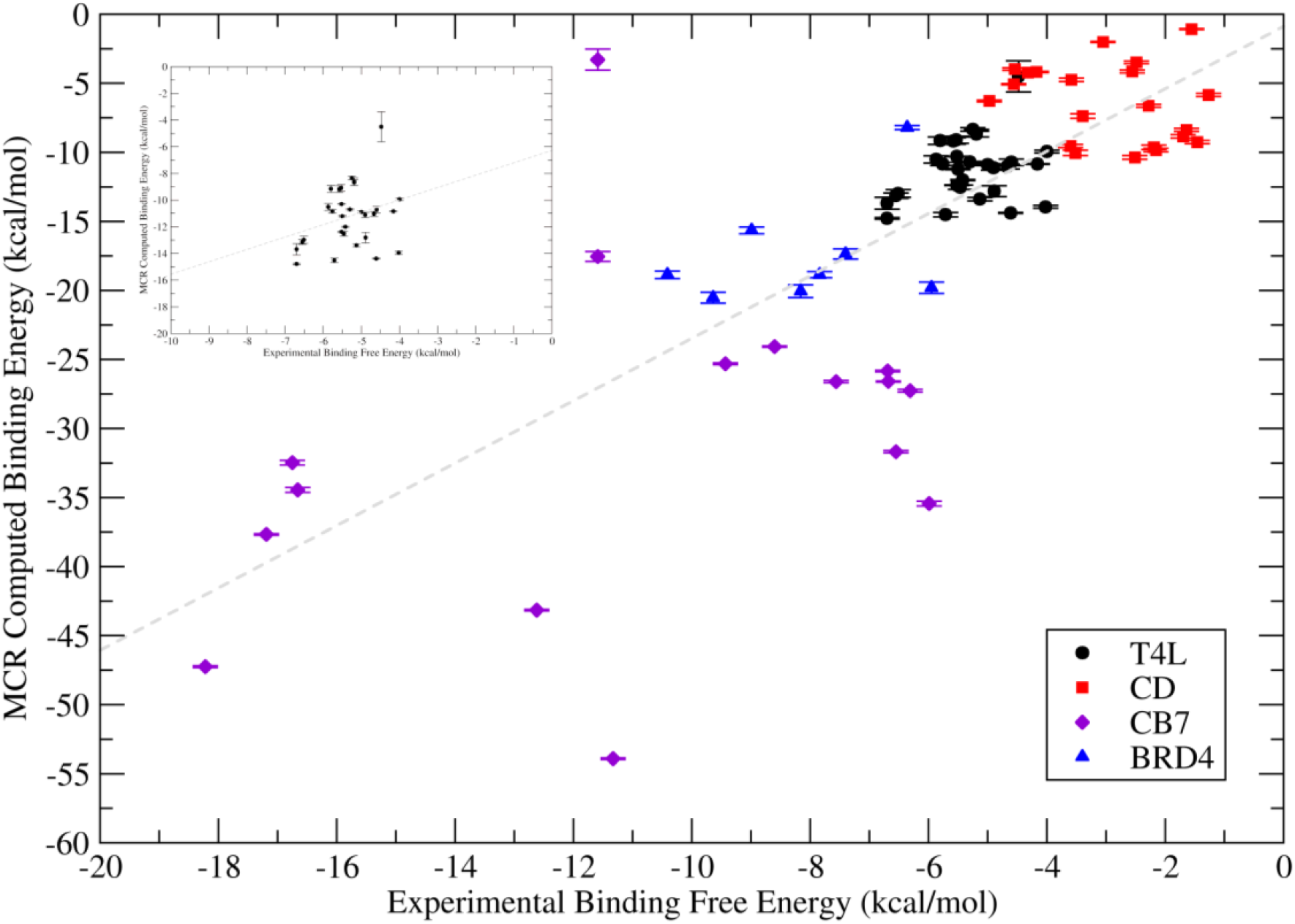
(left) Correlation between the experimental binding free energies (horizontal axis) and binding free energies computed with MCR (vertical axis) for the T4L dataset. (right) Correlation between experimental and computed binding free energies for datasets T4L, CD, CB7, and BRD4 (N=75). The dashed gray line shows the linear regression for the plotted data. The complete dataset is available in the supplementary material.

An inspection of the experimental binding free energies (Table 1) shows that most ligands have binding free energies within the same range. So, given the small range of values found in the experimental results, it would be very difficult for any computational method to make reasonable predictions on this dataset. Methods with a precision of ± 1 kcal/mol, would not distinguish among most of the ligands. So, a more rigorous assessment of the method should include more datasets spanning a wider range of experimental values for the binding free energies.

This issue was further explored by simulating the binding energies in other datasets. Here we choose the datasets CB7, CD, and BRD4. Using the same approaches, the binding energies were computed for these datasets and the results are shown in Figure 4 and Supplementary Material. In this scenario, the experimental binding free energies include values from 0 to -20 kcal/mol and the datasets together account for 75 data points. The correlation between the computed binding free energies and the experimental binding energies, as given by the Pearson correlation coefficient is *r* = 0.76. A bootstrapping analysis assuming a Gaussian error of 0.5 kcal/mol in experimental energies resulted in a correlation coefficient r=0.75±0.01, for N=10^6^ bootstrap iterations. Removing one clear outlier from the CB7 dataset results in a Pearson correlation coefficient of *r* = 0.80. Interestingly, a linear fit between the experimental and computed binding free energies reveals a slope of 2.3 and a linear coefficient of -0.9 kcal/mol, suggesting an important bias of the model towards more negative values, as previously noted.

The data shown in Figure 4 indicates that the MCR method, as applied here, can correctly rank the ligands according to their binding free energy, despite the bias in estimating the absolute binding free energies. At this point, we decided to investigate whether MCR is making better predictions than any ‘end-point’ method, i.e., since the MCR energies are dominated by the low-energy terms at the beginning of the recursion, can average energies computed at low temperatures reach the same performance?

For this purpose, we run Metropolis Monte Carlo simulations at an MC temperature of 300 K for the receptor-ligand complex and the free ligand. The binding energies were computed by taking their differences. In this approach, the conformational entropy was also estimated using a first-order approximation for ligand rotation and translation, as previously proposed by Edholm and Berendsen[16] and by Killian and coworkers[42].

We found that end-point binding energies computed from equilibrium MC simulations resulted in a similar correlation with experimental data when compared to the results obtained with MCR simulations, as shown in Figure 5. This finding suggests that binding energies computed with MCR are mostly based on energy differences sampled at low temperatures.

**Figure 5.**
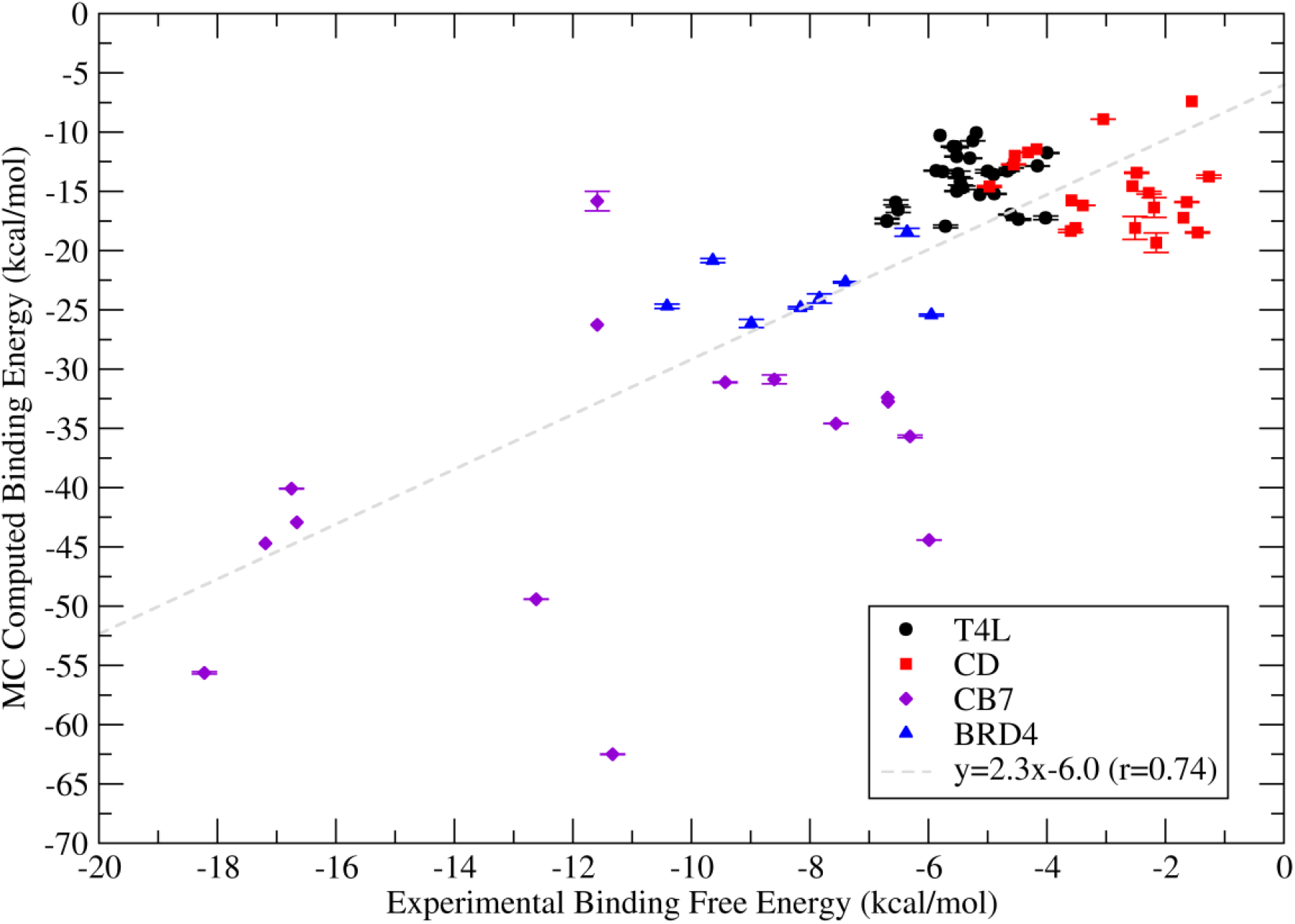
Comparison between binding energies computed with equilibrium Monte Carlo simulations at MC temperature set to T=25K (black points), and MCR simulations (red diamonds). The error bars show the standard deviation for 10 simulations for each receptor-ligand complex. The correlation between computed data (vertical axis) and experimental data (horizontal axis) is r=0.74 for MC and r=0.76 for MCR.

Finally, since the scope of our implementation of the MCR simulation is focused on the refinement of ligand docking scores, we also evaluated the ability of the method to properly rank the ligands after a docking calculation. For this purpose, 13 binders of the L99A mutant of the T4 lysozyme were selected (182L, 183L, 184L, 185L, 186L, 187L, 188L, 4W52, 4W53, 4W55, 4W57, 1NHB, 3DMX) and docked on the crystal structure of the enzyme bound to benzofuran (182L), using the crystallographic ligand (benzofuran) as the *reference* ligand for the docking calculations. The complex obtained in docking calculations was subsequently used in MCR simulations using the same protocol as described above. We found that the docking scores, computed as binding energies as shown in equation (1), have only a very modest correlation with the experimental binding free energies, as shown in Figure 6A, on the top right panel. After an MCR simulation, the energies were re-ranked and showed a better correlation with the experimental data (Figure 6A, top left panel).

**Figure 6.**
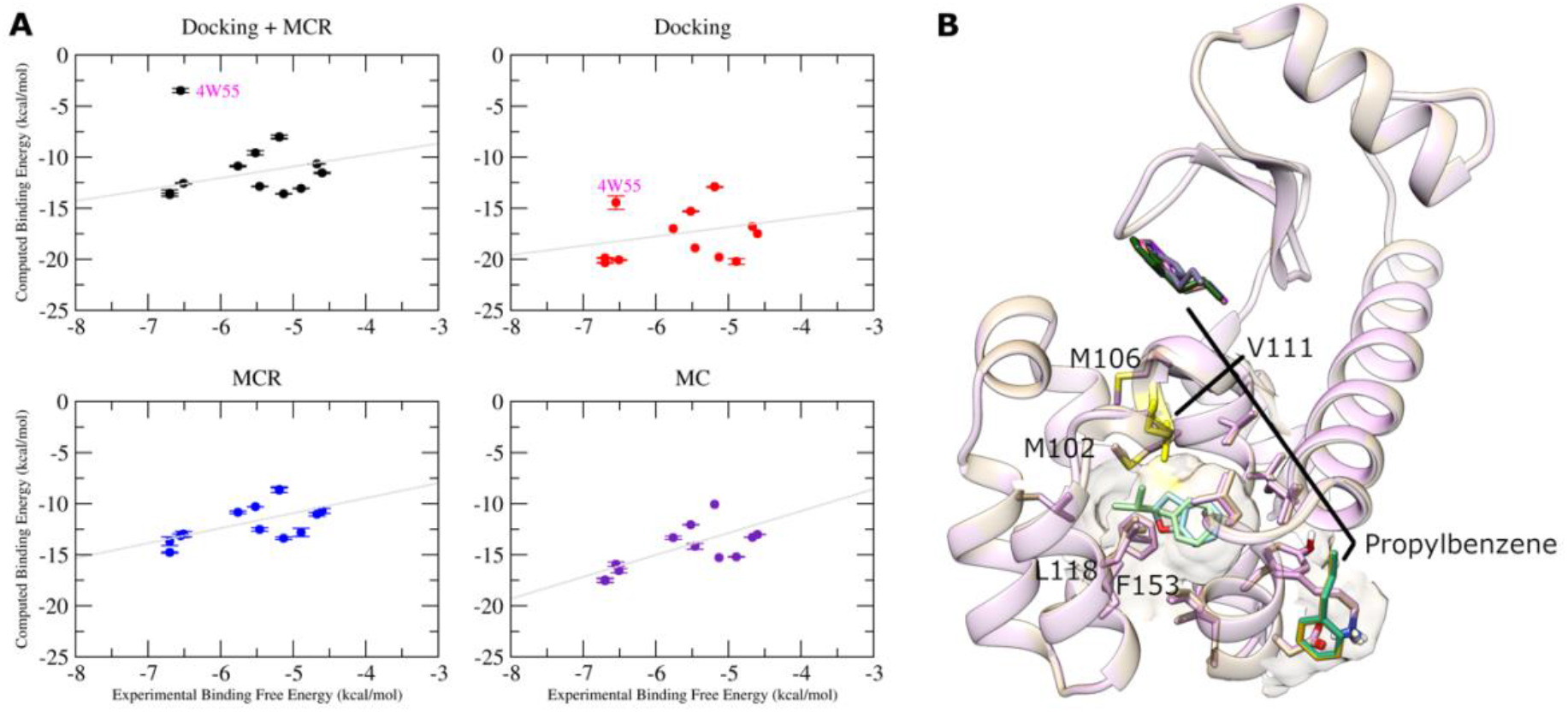
(A) Correlation between experimental binding free energies (horizontal axis) and binding energies computed after ligand docking and MCR calculations (top left), ligand docking only (top right), and MCR or MC calculations using the crystallographic structures of ligand and receptor (bottom left and right, respectively). (B) Superposition of the crystal structures 182L and 4W55 show an important difference in the rotamer for Val111 (shown in yellow sticks). The change in the rotamer impairs the proper docking of propylbenzene in the active site of T4L L99A (PDB ID 182L, originally bound to benzofuran). The crystallographic poses for benzofuran (182L) and propylbenzene (4W55) are shown in blue and green sticks in the active site. The incorrect docking poses are shown as colored sticks on two sites in the enzyme surface.

Interestingly, one complex, 4W55, appears as a clear outlier after MCR calculations. A visual inspection of the coordinates showed that in this complex, a change in the rotamer for Val111 (shown in yellow stick in Figure 6B) was necessary in order to fit the ligand propylbenzene in the nonpolar pocket, as previously noted[7]. Since the receptor was kept rigid, the docking calculations were unable to fit the ligand in the correct pocket. Instead, the ligand was docked on two regions in the surface of the enzyme, as shown in Figure 6B. The incorrectly docked ligand surprisingly showed a binding energy of about -15 kcal/mol in the docking score. However, the MCR calculations corrected the energies to something close to zero, as expected, revealing that the method can distinguish reasonable sites from unreasonable ones.

## 4. Discussion

Here we describe an approach for rapid estimation of the biomolecular interaction by Monte Carlo simulations of the ligand within its binding pocket. The method has a few conveniences: First, the rapid sampling obtained by MC simulations allows its use as a refinement of scores obtained in docking calculations in the context of compound screening. In an ideal scenario, it is plausible to use a docking engine to sample libraries of purchasable compounds containing millions of compounds and pre-select the most likely to bind to a given macromolecular target. Afterward, the pre-selected compounds can be prioritized and ranked using MC simulations, an inexpensive, though reliable method for the estimation of the interaction. Second, the method is extensible and may allow in the future the sampling of a restricted region of the receptor, for example, increasing the reliability of the calculations.

In the approach showed here, we used a rigid receptor and an empirical implicit solvent model. The rigid receptor approach allows the computation of the binding energies using pre-computed grids, as typical in docking calculations [36], significantly contributing to a speed-up in the calculations. On the other hand, relaxation effects due to changes in rotamers close to the active site, for example, are not taken in consideration in our model. The receptor relaxation is somewhat considered in our energy model by using a softcore potential [38]. In the same context, the empirical solvation model used here has the advantage of being compatible with the pre-calculation of terms in the grid approach and was previously shown to be well correlated with experimental solvation free energies [40]. This term was shown to be particularly important to correct the overestimation of the electrostatic term for the binding energies in the Coulomb model.

We showed here that an incorrect pose that showed a reasonable energy score in docking calculations was corrected, in terms of energy score, in subsequent MCR calculations. The results also showed that the MCR calculations are effective even when the calculations are performed for receptor structures that were not originally crystallized with the ligand under investigation, in a scenario closer to real applications. A comparison of MCR calculations of T4L L99A ligands docked against the enzyme structure 182L (Figure 6A, top right) and the crystallographic structure (Figure 6A, bottom left) show similar correlations with experimental data, suggesting that, for this enzyme, even with a rigid receptor, the MCR calculations are efficient in providing a better ranking of the ligands according to their experimental binding energies.

The advantage of MCR relies in the estimation of the partition function using the recursion over the estimation of the amount W(b,T). Other strategies for estimating the partition function include an exhaustive docking [7], for example. In that strategy, a systematic search of the binding modes is generated over the degrees of freedom and then used for a summation to have a partition function. Here, the recursion as proposed by Harold Scheraga provides a means to estimate the partition function over thermodynamic ensembles generated by MC simulations.

In terms of the binding energies computed, the MCR method showed results comparable with end-point calculations taken from equilibrium MC. However, MC simulations are much faster, taking about 6.3 hours on average for each T4L target, while MCR simulations take about 23.6 h for each T4L target, running on a single CPU thread. Although, even with this difference in the required time to simulate, we can point out advantages for MCR over equilibrium MC simulations. First, the MCR method is strongly rooted in thermodynamics. In principle, with better parameters and appropriate sampling, one could end up with free energy values close to experimental data, as shown for simpler systems [24,43]. Second, as we mentioned before, the method may allow the sampling of unbinding coordinates, providing a perspective of the binding funnel for a particular host-guest system.

In terms of the changes in free energy, the ligand binding process can be compared to the protein folding process [22,52,53]: during the binding event, i.e., when moving from the aqueous solvent to a macromolecular binding pocket, a ligand loses its conformational entropy, while gaining interaction energy or enthalpy. The folding funnel model [20], as proposed by Wolynes and Onuchic, has a parallel here for ligand binding, where the correct binding pose should be identified as the global minimum in the funnel. Also, similar to what is found in folding funnels, several local minima are observed in the binding funnel. The roughness of the binding funnel leads to several imprecise results in calculations based on finding (local) minima, such as in docking calculations, for example, resulting in a poor correlation between docking scores and experimental data. The MCR method, when applied to ligand binding, as shown in this work, can be seen as a computational method to sample low-energy conformations (bottom of the funnel), as well as high-energy conformations, potentially providing an assessment of the accessible conformations in the (un)binding funnel. In this context, additional investigation is underway to evaluate whether unbinding events sampled in MCR correlate with experimental ligand unbinding kinetic data, for example.

A representative ligand-receptor view for this binding funnel can be seen in Figure 7 for the complexes 185L and 4W57, from the T4L dataset. The figure shows that conformations with low RMSD values (as compared to the crystal structure conformation) are observed for low-energy conformations. As the energy increases, the observed RMSD values also increase, and, at RMSD values close to 10 Å, the ligand is dissociated from its receptor. After unbinding, the ligand samples a wide range of conformations with RMSD values varying between 10 to 30 Å. The conformations with RMSD in the range 5-10 Å typically show a high energy barrier that the ligand faces before reaching the low-energy bound state. The comparison of the landscapes shown in Figure 7 for indole (185L) and N-butylbenzene (4W57) binding to T4L L99A also reveals that indole has a broader energy minimum, locate at RMSD 0-5Å, while the energy minima for N-butylbenzene is more well-defined in two clusters of conformations at RMSD 0-2.5Å and about 5 Å. Interestingly, the experimental data provided by Morton and Matthews [46] show that the entropic component for the binding free energy is increased for indole, as compared to N-butylbenze. In indole binding, a Δ*G*_*bind*_ = −4.89 kcal/mol is observed with an entropic component −*T*Δ*S*_*bind*_ = 6.34 kcal/mol, while for N-butylbenze, a Δ*G*_*bind*_ = −6.7 kcal/mol is observed with an entropic component −*T*Δ*S*_*bind*_ = 1.36 kcal/mol. This comparison between computed and experimental data suggests that the Monte Carlo strategy as applied in this work is properly sampling interesting properties associated with the binding thermodynamics of small ligands in the model system T4L.

**Figure 7.**
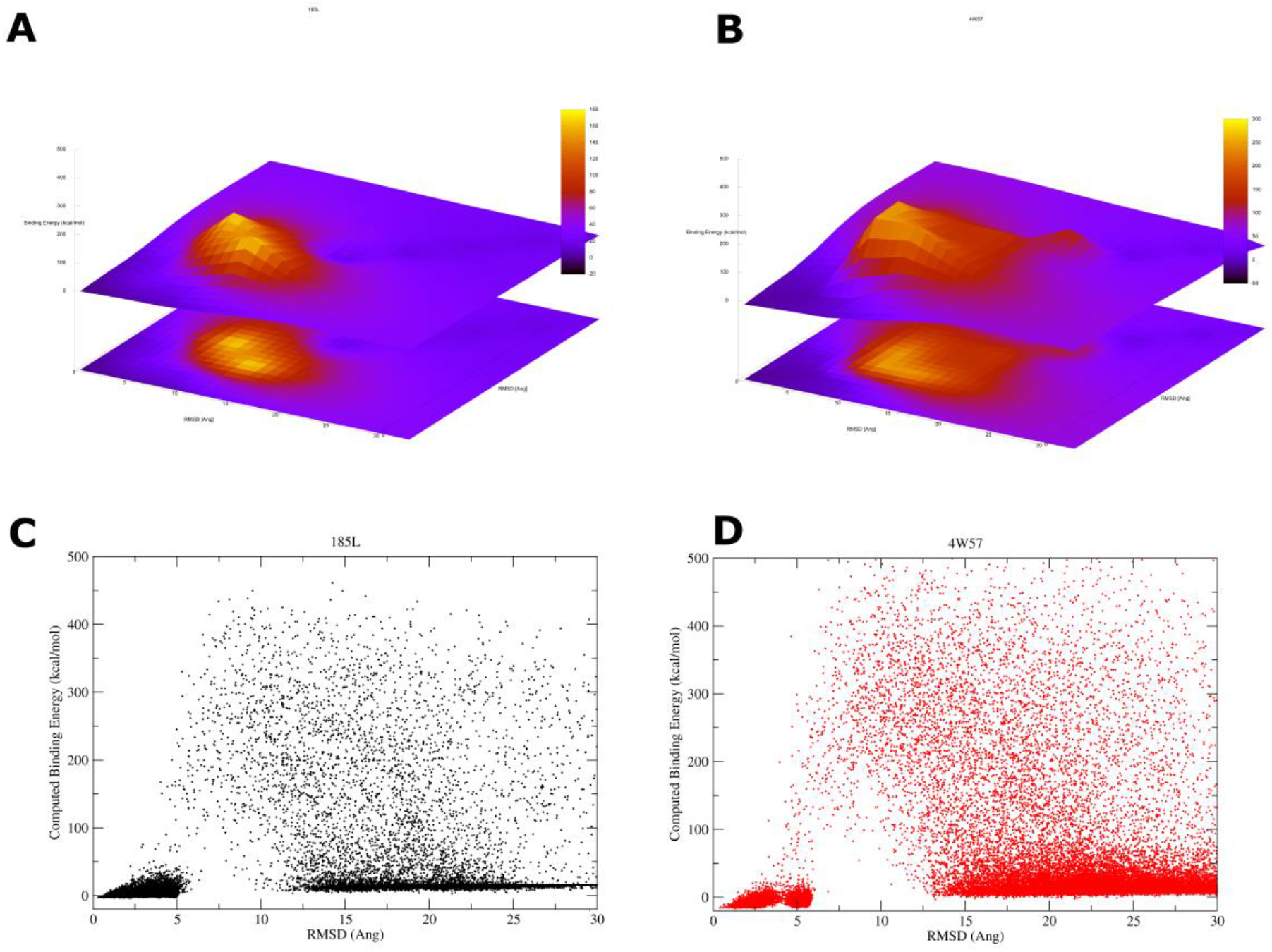
Graphical representation of the RMSD observed for the ligand versus the total energy (E_t_). Panels (A) and (B) show the ligand RMSD in the x-z plane versus Energy in the y-axis for the complexes 185L and 4W57, respectively. Panels (C) and (D) show the same data in a two-dimension plot. Note that indole (185L) shows a broader energy minimum when compared to N-butylbenzene.

When compared to other methods for the estimation of the binding energy, the MCR method shows promising results. Steinbrecher and coworkers evaluated the FEP+ approach for FEP calculations using a dataset of 12 T4L L99A ligands [8]. The authors found a correlation coefficient of *r*=0.87 for FEP+ with an RMSE of 0.7 kcal/mol. Interestingly, the authors also compared their method with Glide SP docking scores (*r*=0.57) and with MM-GBSA (*r*=0.55). The calculations with MCR showed a correlation coefficient of *r*=0.59 for N=13 T4L L99A ligands or *r*=0.67 for equilibrium MC calculations (Supplementary Material). The correlations obtained given the assumptions made show that the method can be considered as an intermediate approach, cheaper than alchemical methods, but more precise than single point calculations, such as docking scores.

On the other hand, the MCR simulation is unique in terms of the sampling strategy it adopts. For example, the BEDAM approach, Binding Energy Distribution Analysis Method, proposed by Gallicchio, Levy, and coworkers [54] estimates the interaction energies and their probabilities from Umbrella sampling molecular dynamics with Hamiltonian replica exchanges. The authors showed very interesting results for eight T4L L99A ligands and, their results were better in ranking the ligands properly than in the estimation of the experimental binding free energies. The blurring approach, proposed by Merz and coworkers [6], and the exhaustive docking approach as proposed by Purisima and Hogues [7] are both based on the sampling of low-energy conformations close to the global minimum and an estimation of the partition function with limited sampling. In both cases, promising results were shown in terms of the correlation of the computed binding energies and the experimental binding free energies, but with overestimation in the computed energies. The MCR approach is distinct in the sense that the sampling is obtained in a non-equilibrium dataset, simulated in increasing effective temperatures, providing a sampling of the energy minimum as well as high-energy states. Another distinctive feature is the lack of direct estimation of the configurational integrals, making the estimation of the free energies easier.

Finally, one important point is whether the MCR calculations can readily generalize to more diverse protein-ligand complexes. Preliminary tests with the CASF-2016 dataset [55] indicate that MCR can distinguish strong binders from weaker ones (Supplementary Figure 4), suggesting that the method is generalizable for more diverse and intricate protein-ligand complexes.

## 5. Conclusions

In conclusion, the data provided here show the conceptually simple approach for the determination of the binding free energy by combining an MC ensemble average at increasing effective temperatures and free energy estimation using the Monte Carlo Recursion method. The method allows a computationally cheap refinement of the binding poses and binding energies obtained by docking calculations, potentially making the structure-based design of drugs more assertive. The MCR calculations with an extended dataset containing T4L, CD and CB7 showed a correlation with experimental data (*r*=0.76) which is better than the correlation previously observed for T4L complexes using MM-GBSA (*r*=0.55) or docking scores (*r*=0.57). The direct evaluation of this thermodynamic quantity allows a more precise ranking of screened ligands in docking campaigns at an affordable computational cost, even for small and in-house computer clusters. The codes developed for this analysis are publicly available on GitHub as a part of the `LiBELa`/`MCLiBELa` project (https://github.com/alessandronascimento/LiBELa).

## Acknowledgments

The authors thank the funding agencies FAPESP for the financial support through grants 2014/06565-2, 2017/18173-0, 2020/03983-9, 2010/15376-8, 2015/26722-8, as well as for CNPq, through grants 485950/2013-8 and 302992/2021-9. JVSC also thanks FAPESP for the Master fellowship 2015/01709-9 and 2014/01751-2. This study was financed in part by the Coordenação de Aperfeiçoamento de Pessoal de Nível Superior - Brasil (CAPES) - Finance Code 001. We also thank Heloisa Muniz, Camila Tanimoto Rodrigues, and Milton T. Sonoda (*in memorian*) for the very fruitful discussions.

## Table of Contents/Abstract Graphics

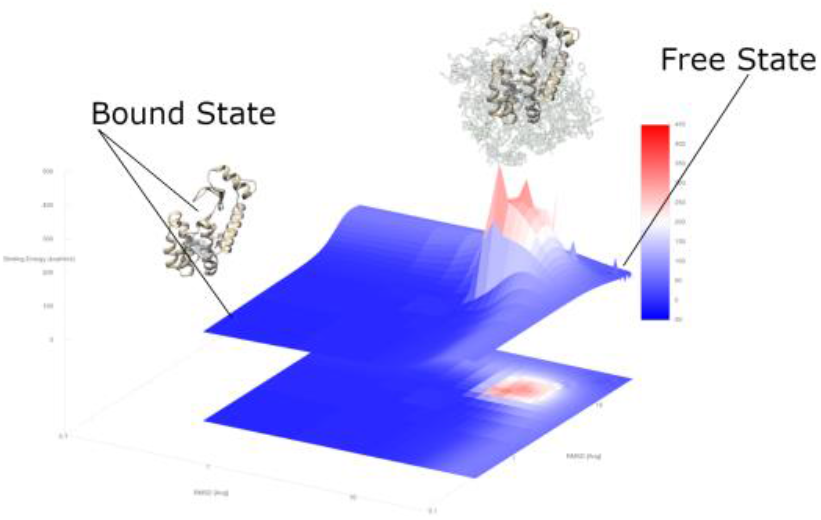

