## Supplementary Material for "Ligand Binding Free Energy Evaluation by Monte Carlo Recursion"

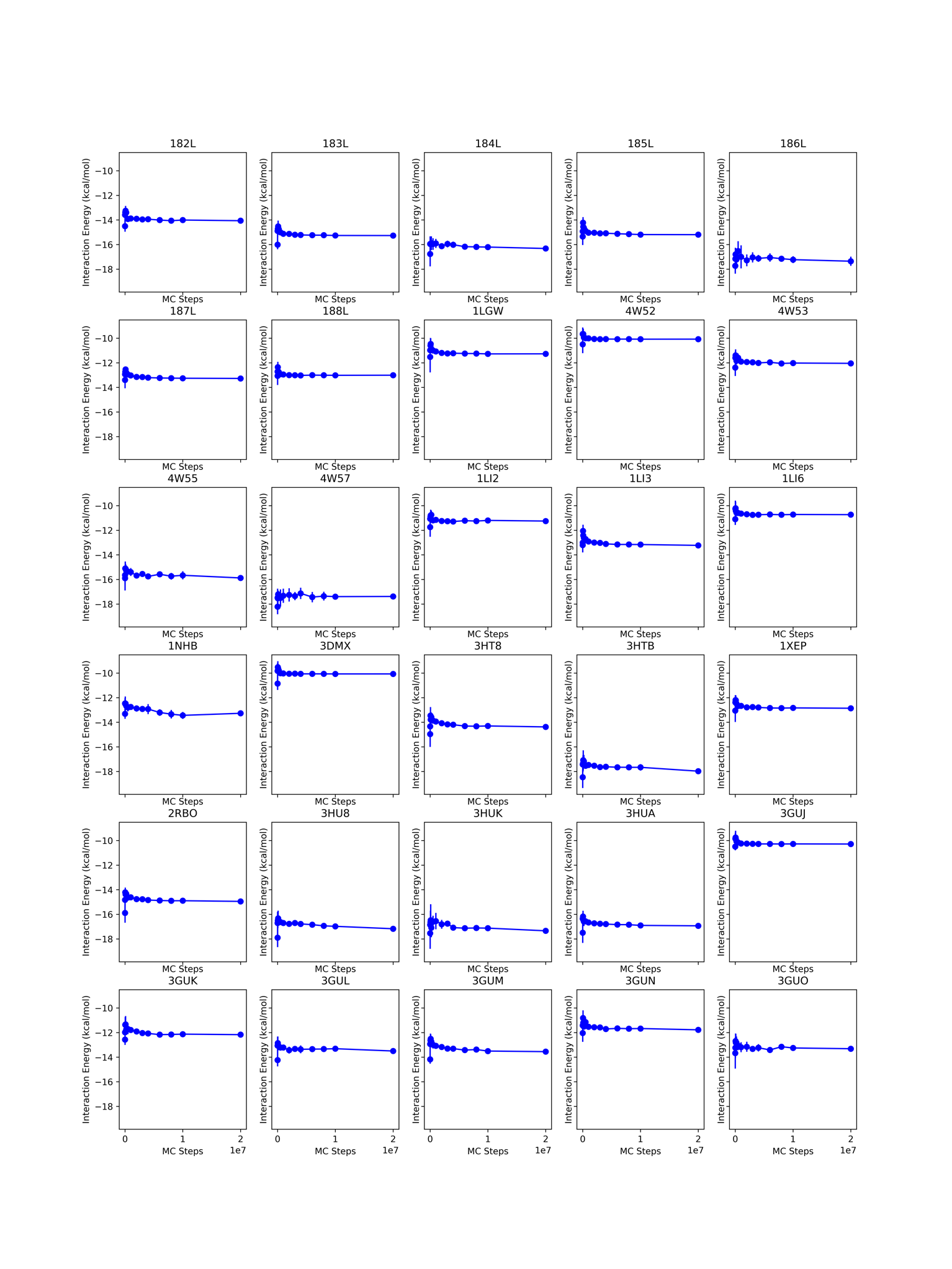

Supplementary Figure 1. Convergence of Binding Energies in equilibrium MC simulations. The vertical axis show the Binding Energy computed as a sum of the $E_{i}=\left\langle\Delta U \right\rangle-T\Delta S$. The entropy is estimated using the first-order approach and binning of each degree of freedom for the ligand (rotational, translational and torsional) and computing the entropy of each degree of freedom as a Shannon entropy $\frac{S_{i}}{k}=-p_{i}\ln p_{i}$.

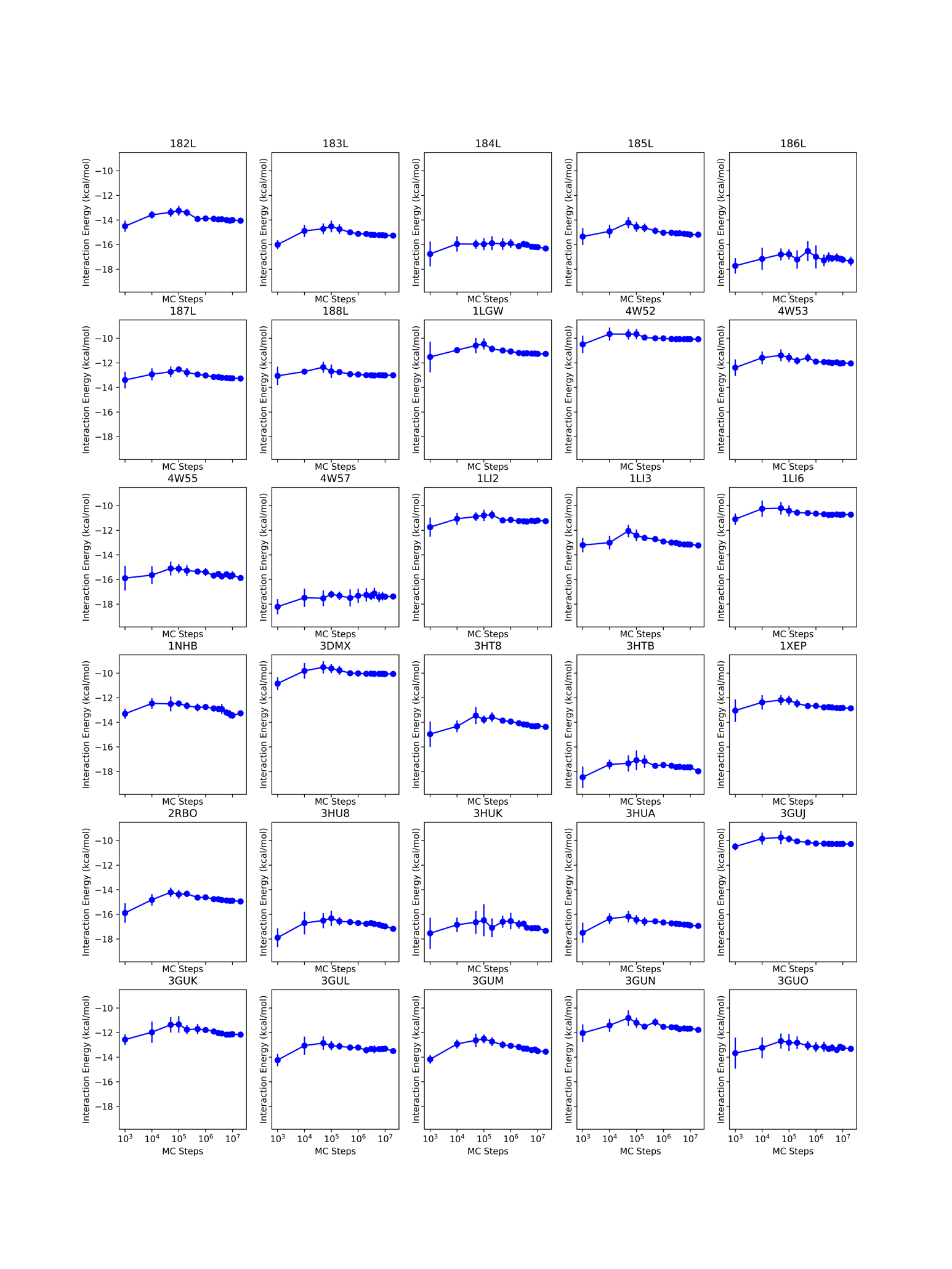

Supplementary Figure 2. Same data as shown in Figure 1 above. Here the MC steps (horizontal axis) are shown in logarithmic scale.

Supplementary Table 1. Experimental data and computed energies after Docking and MCR simulations (N=10) with T4L L99A ligands. All the docking calculations used the enzyme structure available in the PDB ID 182L and the benzofuran as the reference ligand for docking.

|  |  | **Experimental Data** | | | | **Computed data** | |
| --- | --- | --- | --- | --- | --- | --- | --- |
| **ID** | **Ligand** | **ΔG**  **(kcal/mol)** | **ΔH**  **(kcal/mol)** | **ΔH error**  **(kcal/mol)** | **-TΔS**  **(kcal/mol)** | **E_b_**  **(kcal/mol)** | **SD**  **(kcal/mol)** |
| 182L | Benzofuran | -5.46 | -8.04 | 0.44 | 2.58 | -12.87 | 0.03 |
| 183L | Indene | -5.13 | -8.31 | 0.48 | 3.18 | -13.60 | 0.02 |
| 184L | isobutylbenzene | -6.51 | -7.09 | 0.35 | 0.58 | -12.56 | 0.06 |
| 185L | Indole | -4.89 | -11.23 | 0.94 | 6.34 | -13.07 | 0.03 |
| 186L | N-butylbenzene | -6.7 | -8.06 | 0.98 | 1.36 | -13.67 | 0.1 |
| 187L | Para-xylene | -4.67 | -6.97 | 0.98 | 2.3 | -10.67 | 0.02 |
| 188L | O-xylene | -4.6 | -8.45 | 0.96 | 3.85 | -11.54 | 0.07 |
| 4W52 | Benzene | -5.19 | -6.32 | 0.37 | 1.13 | -8.0 | 0.2 |
| 4W53 | Toluene | -5.52 | -6.53 | 0.73 | 1.01 | -9.6 | 0.2 |
| 4W55 | Propylbenzene | -6.55 | -9.97 | 0.05 | 3.42 | -3.5 | 0.2 |
| 4W57 | N-butylbenzene | -6.7 | -8.06 | 0.98 | 1.36 | -13.5 | 0.3 |
| 1NHB | Phenylethane | -5.76 | -6.76 | -- | 1 | -10.88 | 0.07 |
| 3DMX | Benzene | -5.19 | -6.32 | 0.37 | 1.13 | -8.1 | 0.2 |

Supplementary Table 2. Experimental data and computed energies after Docking calculations (N=10) with T4L L99A ligands. All the docking calculations used the enzyme structure available in the PDB ID 182L and the benzofuran as the reference ligand for docking.

|  |  | **Experimental Data** | | | **Docking Data** | |
| --- | --- | --- | --- | --- | --- | --- |
| **Target** | **Ligand** | **ΔG (kcal/mol)** | **ΔH**  **(kcal/mol)** | **-TΔS**  **(kcal/mol)** | **Docking Energy**  **(kcal/mol)** | **SD** |
| 182L | Benzofuran | -5.46 | -8.04 | 2.58 | -18.9 | 0 |
| 183L | Indene | -5.13 | -8.31 | 3.18 | -19.8 | 0 |
| 184L | isobutylbenzene | -6.51 | -7.09 | 0.58 | -20.07 | 0.05 |
| 185L | Indole | -4.89 | -11.23 | 6.34 | -20.2 | 0.3 |
| 186L | N-butylbenzene | -6.7 | -8.06 | 1.36 | -20.35 | 0.05 |
| 187L | Para-xylene | -4.67 | -6.97 | 2.3 | -16.8 | 0 |
| 188L | O-xylene | -4.6 | -8.45 | 3.85 | -17.5 | 0 |
| 4W52 | Benzene | -5.19 | -6.32 | 1.13 | -12.92 | 0.06 |
| 4W53 | Toluene | -5.52 | -6.53 | 1.01 | -15.31 | 0.06 |
| 4W55 | Propylbenzene | -6.55 | -9.97 | 3.42 | -14.45 | 0.7 |
| 4W57 | N-butylbenzene | -6.7 | -8.06 | 1.36 | -19.87 | 0.05 |
| 1NHB | Phenylethane | -5.76 | -6.76 | 1 | -17.0 | 0 |
| 3DMX | Benzene | -5.19 | -6.32 | 1.13 | -12.89 | 0.03 |

Supplementary Table 3. Experimental data and computed energies with MCR simulations. N=10 and the associated error is computed as the standard deviation among the 10 replicates.

| **Set** | **Target** | **Exptl ΔG** | **Calc E** | **Err(E)** |
| --- | --- | --- | --- | --- |
| T4L | 182L | -5.46 | -12.52 | 0.16 |
|  | 183L | -5.13 | -13.39 | 0.12 |
|  | 184L | -6.51 | -12.98 | 0.28 |
|  | 185L | -4.89 | -12.83 | 0.40 |
|  | 186L | -6.7 | -14.79 | 0.06 |
|  | 187L | -4.67 | -11.03 | 0.17 |
|  | 188L | -4.6 | -10.72 | 0.24 |
|  | 1LGW | -5.53 | -9.09 | 0.25 |
|  | 4W52 | -5.19 | -8.68 | 0.21 |
|  | 4W53 | -5.52 | -10.30 | 0.01 |
|  | 4W55 | -6.55 | -13.14 | 0.19 |
|  | 4W57 | -6.7 | -13.69 | 0.43 |
|  | 1LI2 | -5.58 | -9.18 | 0.24 |
|  | 1LI3 | -5.87 | -10.51 | 0.26 |
|  | 1LI6 | -5.25 | -8.35 | 0.12 |
|  | 1NHB | -5.76 | -10.84 | 0.12 |
|  | 3DMX | -5.19 | -8.62 | 0.28 |
|  | 3HT8 | -5.42 | -12.01 | 0.04 |
|  | 3HTB | -5.71 | -14.52 | 0.16 |
|  | 1XEP | -4.16 | -10.84 | 0.02 |
|  | 2RBO | -5.52 | -12.38 | 0.03 |
|  | 3HU8 | -4.02 | -13.95 | 0.10 |
|  | 3HUK | -4.48 | -4.51 | 1.13 |
|  | 3HUA | -4.61 | -14.39 | 0.03 |
|  | 3GUJ | -5.8 | -9.16 | 0.27 |
|  | 3GUK | -5.3 | -10.70 | 0.05 |
|  | 3GUL | -5.5 | -11.21 | 0.05 |
|  | 3GUM | -4.9 | -11.11 | 0.18 |
|  | 3GUN | -4 | -9.94 | 0.10 |
|  | 3GUO | -5 | -10.89 | 0.07 |
| CD | 2_1 | -2.486 | -3.49 | 0.09 |
|  | 2_2 | -3.585 | -4.74 | 0.12 |
|  | 2_3 | -3.05 | -2.01 | 0.04 |
|  | 2_4 | -4.541 | -3.98 | 0.09 |
|  | 2_5 | -4.56 | -5.06 | 0.04 |
|  | 2_6 | -1.267 | -5.85 | 0.12 |
|  | 2_7 | -3.394 | -7.38 | 0.18 |
|  | 2_8 | -1.64 | -8.38 | 0.11 |
|  | 2_9 | -1.697 | -8.85 | 0.12 |
|  | 2_10 | -2.192 | -9.64 | 0.16 |
|  | 2_11 | -2.512 | -10.37 | 0.13 |
|  | 2_12 | -3.599 | -9.55 | 0.11 |
|  | 2_s13 | -2.557 | -4.14 | 0.10 |
|  | 2_s14 | -1.554 | -1.08 | 0.04 |
|  | 2_s15 | -4.175 | -4.18 | 0.05 |
|  | 2_s16 | -4.319 | -4.23 | 0.09 |
|  | 2_s17 | -4.971 | -6.28 | 0.07 |
|  | 2_s18 | -2.28 | -6.64 | 0.10 |
|  | 2_s19 | -1.458 | -9.26 | 0.12 |
|  | 2_s20 | -2.156 | -9.84 | 0.13 |
|  | 2_s21 | -3.518 | -10.05 | 0.20 |
| CB7 | 1_17 | -11.33 | -53.91 | 0.06 |
|  | 1_18b | -11.59 | -3.29 | 0.76 |
|  | 1_18 | -11.59 | -17.55 | 0.36 |
|  | 1_1 | -5.99 | -35.44 | 0.18 |
|  | 1_22 | -16.66 | -34.45 | 0.17 |
|  | 1_23 | -17.19 | -37.67 | 0.06 |
|  | 1_24 | -16.75 | -32.47 | 0.17 |
|  | 1_3 | -6.55 | -31.68 | 0.08 |
|  | 1_5 | -18.22 | -47.24 | 0.07 |
|  | 2_20 | -12.62 | -43.15 | 0.07 |
|  | 2_2 | -6.31 | -27.26 | 0.10 |
|  | 2_4 | -6.68 | -26.59 | 0.05 |
|  | 2_5 | -6.69 | -25.84 | 0.06 |
|  | 2_7 | -7.56 | -26.61 | 0.08 |
|  | 2_8 | -8.6 | -24.07 | 0.05 |
|  | 2_9 | -9.43 | -25.32 | 0.07 |
| BRD4 | 3 | -5.95 | -19.81 | 0.41 |
|  | 4 | -6.36 | -8.21 | 0.14 |
|  | 5 | -7.4 | -17.36 | 0.36 |
|  | 6 | -7.84 | -18.86 | 0.22 |
|  | 7 | -8.16 | -20.06 | 0.46 |
|  | 8 | -8.99 | -15.66 | 0.24 |
|  | 9 | -9.64 | -20.52 | 0.40 |
|  | 10 | -10.41 | -18.87 | 0.27 |

Supplementary Table 4. Experimental data and computed energies with equilibrium MC simulations (T=300 K). N=10 and the associated error is computed as the standard deviation among the 10 replicates.

| **Set** | **Target** | **Exptl ΔG** | **Calc E** | **Err(E)** |
| --- | --- | --- | --- | --- |
| T4L | 182L | -5.46 | -14.18 | 0.28 |
|  | 183L | -5.13 | -15.28 | 0.00 |
|  | 184L | -6.51 | -16.56 | 0.22 |
|  | 185L | -4.89 | -15.21 | 0.04 |
|  | 186L | -6.7 | -17.48 | 0.22 |
|  | 187L | -4.67 | -13.29 | 0.01 |
|  | 188L | -4.6 | -13.01 | 0.02 |
|  | 1LGW | -5.53 | -11.25 | 0.07 |
|  | 4W52 | -5.19 | -10.08 | 0.00 |
|  | 4W53 | -5.52 | -12.06 | 0.03 |
|  | 4W55 | -6.55 | -15.93 | 0.20 |
|  | 4W57 | -6.7 | -17.55 | 0.20 |
|  | 1LI2 | -5.58 | -11.23 | 0.05 |
|  | 1LI3 | -5.87 | -13.26 | 0.04 |
|  | 1LI6 | -5.25 | -10.74 | 0.01 |
|  | 1NHB | -5.76 | -13.34 | 0.15 |
|  | 3DMX | -5.19 | -10.07 | 0.00 |
|  | 3HT8 | -5.42 | -14.70 | 0.15 |
|  | 3HTB | -5.71 | -17.96 | 0.11 |
|  | 1XEP | -4.16 | -12.86 | 0.02 |
|  | 2RBO | -5.52 | -14.98 | 0.04 |
|  | 3HU8 | -4.02 | -17.24 | 0.15 |
|  | 3HUK | -4.48 | -17.37 | 0.08 |
|  | 3HUA | -4.61 | -16.96 | 0.03 |
|  | 3GUJ | -5.8 | -10.29 | 0.00 |
|  | 3GUK | -5.3 | -12.20 | 0.03 |
|  | 3GUL | -5.5 | -13.52 | 0.24 |
|  | 3GUM | -4.9 | -13.58 | 0.07 |
|  | 3GUN | -4 | -11.76 | 0.05 |
|  | 3GUO | -5 | -13.31 | 0.13 |
| CD | 2_1 | -2.486 | -13.43 | 0.07 |
|  | 2_2 | -3.585 | -15.77 | 0.00 |
|  | 2_3 | -3.05 | -8.92 | 0.01 |
|  | 2_4 | -4.541 | -12.00 | 0.00 |
|  | 2_5 | -4.56 | -12.73 | 0.04 |
|  | 2_6 | -1.267 | -13.76 | 0.13 |
|  | 2_7 | -3.394 | -16.18 | 0.01 |
|  | 2_8 | -1.64 | -15.90 | 0.04 |
|  | 2_9 | -1.697 | -17.22 | 0.00 |
|  | 2_10 | -2.192 | -16.36 | 0.83 |
|  | 2_11 | -2.512 | -18.08 | 0.97 |
|  | 2_12 | -3.599 | -18.34 | 0.13 |
|  | 2_s13 | -2.557 | -14.55 | 0.00 |
|  | 2_s14 | -1.554 | -7.41 | 0.00 |
|  | 2_s15 | -4.175 | -11.44 | 0.00 |
|  | 2_s16 | -4.319 | -11.71 | 0.03 |
|  | 2_s17 | -4.971 | -14.59 | 0.08 |
|  | 2_s18 | -2.28 | -15.12 | 0.11 |
|  | 2_s19 | -1.458 | -18.47 | 0.06 |
|  | 2_s20 | -2.156 | -19.33 | 0.82 |
|  | 2_s21 | -3.518 | -18.09 | 0.00 |
| CB7 | 1_17 | -11.33 | -62.50 | 0.03 |
|  | 1_18b | -11.59 | -15.82 | 0.82 |
|  | 1_18 | -11.59 | -26.26 | 0.00 |
|  | 1_1 | -5.99 | -44.42 | 0.01 |
|  | 1_22 | -16.66 | -42.93 | 0.00 |
|  | 1_23 | -17.19 | -44.70 | 0.00 |
|  | 1_24 | -16.75 | -40.08 | 0.03 |
|  | 1_5 | -18.22 | -55.63 | 0.09 |
|  | 2_20 | -12.62 | -49.42 | 0.02 |
|  | 2_2 | -6.31 | -35.68 | 0.11 |
|  | 2_4 | -6.68 | -32.75 | 0.00 |
|  | 2_5 | -6.69 | -32.39 | 0.00 |
|  | 2_7 | -7.56 | -34.59 | 0.02 |
|  | 2_8 | -8.6 | -30.87 | 0.37 |
|  | 2_9 | -9.43 | -31.11 | 0.04 |
| BRD4 | 3 | -5.95 | -25.43 | 0.09 |
|  | 4 | -6.36 | -18.45 | 0.34 |
|  | 5 | -7.4 | -22.68 | 0.08 |
|  | 6 | -7.84 | -24.04 | 0.40 |
|  | 7 | -8.16 | -24.83 | 0.10 |
|  | 8 | -8.99 | -26.14 | 0.34 |
|  | 9 | -9.64 | -20.84 | 0.18 |
|  | 10 | -10.41 | -24.69 | 0.18 |

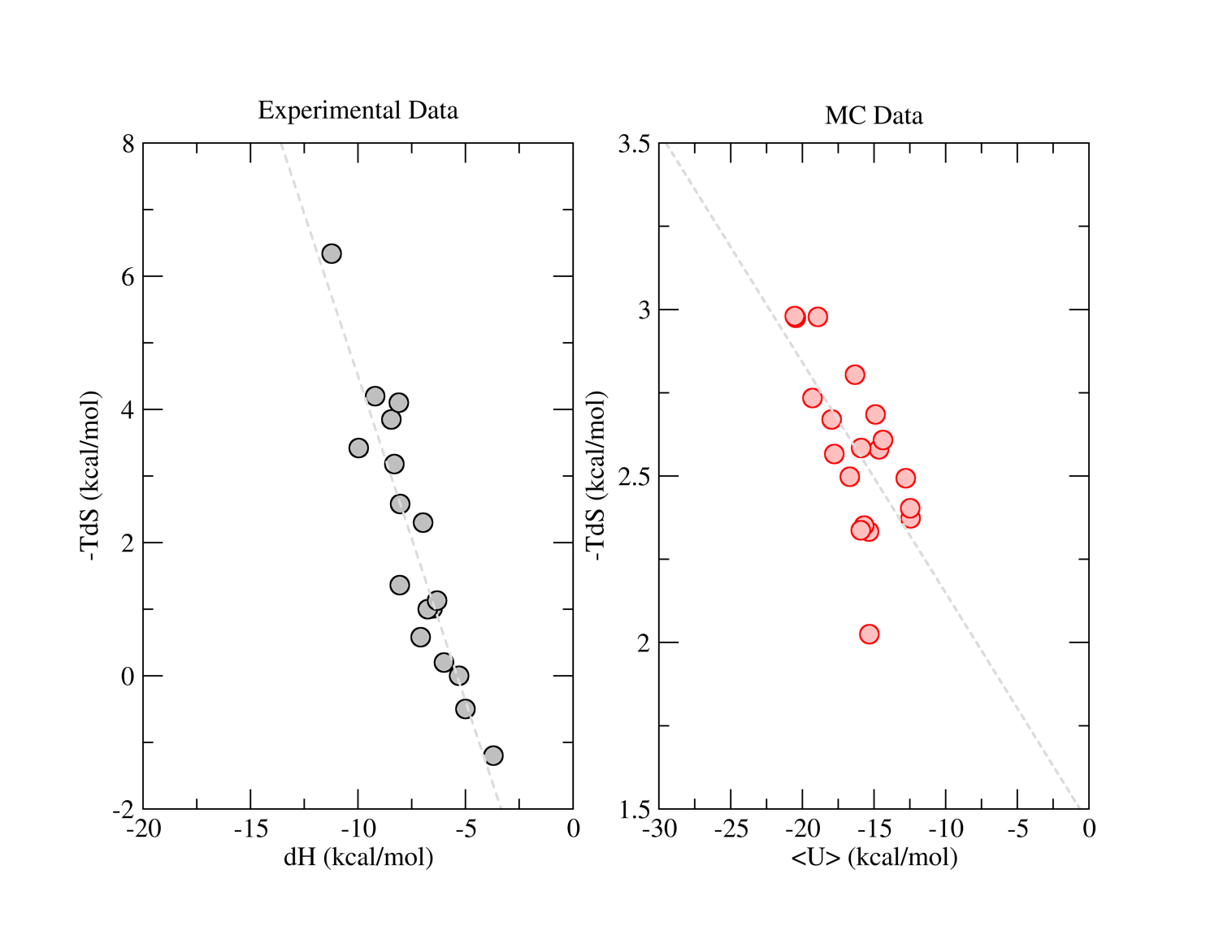

Supplementary Figure 3. Thermodynamic Profile of T4 ligands. (left) Experimental binding entropies (-TdS) are ploted against experimental binding enthalpies, revealing a linear correlation. (right) Entlhalpies and entropies computed from equilibrium MC simulations show a similar trend, although overestimating the enthalpies and underestimating the entropies. Note that the computed entropies are configurational entropies, i.e., entropic contributions of solvation, for example, are not included in the calculations.

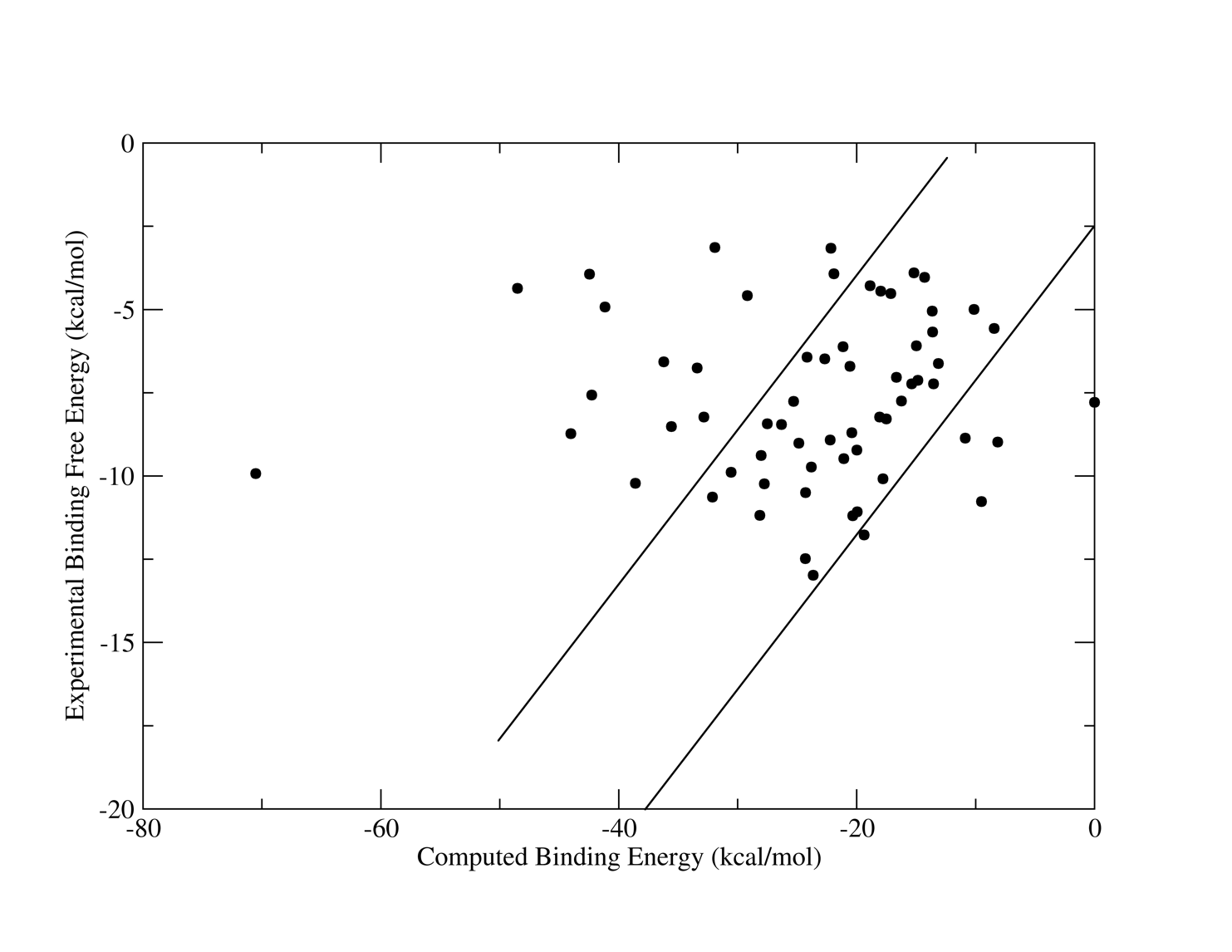

Supplementary Figure 4. Correlation between experimental binding free energies (vertical axis) and binding energies computed with MCR (horizontal axis) for the CASF-2016 dataset. The CASF dataset was filtered to include only complexes with binding experimental data (Kd). 62 protein-ligand complexes were used in the calculations as provided by PDBBind website. The parameters for the calculations were kept as the same as used for the T4L dataset. The lines show a set of complexes with a good correlation between computed and experimental data.

Typical input files used in MCR simulations.

```
# mode

mode					mcr
dock_parallel          		yes
parallel_jobs			1

#
# input files
#

rec_mol2                	../../../../mol2/182L_DOCKPREP.mol2.gz
lig_mol2                	../../../../mol2/182L_LIG_DOCKPREP.mol2.gz
reflig_mol2				../../../../mol2/182L_LIG_DOCKPREP.mol2.gz
multifile               	multimol.dat

#
# force field parameters
#

deltaij				2.5
deltaij_es				2.5
mol2_aa				no
search_box				45.0 45.0 45.0
scoring_function        	0
use_grids				yes
grid_spacing            	0.4
grid_box                	50.0    50.0    50.0
load_grids				../McGrid
solvation_alpha			0.1
dielectric_model			r
diel					2.0
ligand_energy_model		GAFF
atomic_model_ff			AMBER
LJ_sigma				1.2

#
# PBSA Parameter
#
use_pbsa                	no
use_delphi				no

#
# Optimization
#

timeout                 	120
minimization_tolerance  	1.0e-10
minimization_delta      	1.0e-5
dock_min_tol           	 	1.0e-10
minimization_timeout   		120
sort_by_energy			yes
overlay_optimizer       	ln_auglag
energy_optimizer      		direct
ignore_h                	no
deal                    	no
elec_scale				1.0
vdw_scale				1.0

#
# MC Options
#
nsteps				20000000
temperature				300.0
cushion				0.5
max_atom_displacement		0.035
rotation_step			1.25
torsion_step			1.25
sample_torsions			yes
mc_full_flex			no
compute_rotation_entropy	no
equilibration_steps		1000000
ligand_simulation			yes
mc_stride				50000
seed 					12477
#entropy_rotation_bins		720
#entropy_translation_bins	20

#
# SA Options
#
sa_start_temp			100.0
sa_steps				10000
sa_mu_t				1.1

#
# MCR Options
#

mcr_size             	   	12
mcr_coefficients      	  	1.5 1.5 2.0 2.0 2.0 4.0 8.0 16.0 32.0 64.0 128.0 256.0

#
# output
#

output_prefix			182L_SF0_1
write_mol2				yes

#
# flexible ligands
#

generate_conformers		no
number_of_conformers		10
conformers_to_rank		1
conformer_min_steps		1000
```
